# NeuroTD infers time varying delays in neural activities by adaptive sliding window alignment

**DOI:** 10.1101/2024.10.28.620662

**Authors:** Xiang Huang, Noah Cohen Kalafut, Sayali Anil Alatkar, Athan Z. Li, Qiping Dong, Qiang Chang, Daifeng Wang

## Abstract

Studying the temporal dynamics of neural activities is essential for understanding how neurons function. These dynamics often involve temporal delays between neurons that vary over time, revealing both their functions and how they interact within circuits. Recent techniques such as Neuropixels, depth electrodes, and Patch-seq enable time-series recordings of neural activity at various scales, ranging from single neurons to large populations. However, inferring such time-varying delays remains challenging due to noise, high sampling rates, and complex temporal patterns. To address these challenges, we developed NeuroTD, a novel computational approach based on sliding windows to align time-series datasets and infer time-varying delays. Particularly, NeuroTD integrates adaptive window-size tuning to obtain optimal and robust delay estimates. We first benchmarked NeuroTD in simulation studies, demonstrating its robustness and outperformance. Then we applied it to two emerging real-world datasets: (i) intracranial multi-channel electrophysiological recordings from depth electrodes across medial temporal lobe regions in humans, showing that hippocampal signals recorded via depth electrodes exhibited consistently significantly longer time delays than other regions during working memory tasks, and (ii) Patch-seq data in the mouse motor cortex, revealing intrinsic electrophysiological time-delays of excitatory neurons correlated with gene expression and highlighting pathways related to ion transport and neuronal excitability. Finally, NeuroTD is open-source and available at https://github.com/daifengwanglab/NeuroTD for general use.

## 1 Introduction

Understanding the temporal dynamics of neural activities is essential for decoding neural functions and connectivity. These dynamics often involve specific time delays, where the sign of the delay indicates whether one signal leads or lags another, providing insights into interactions within neural circuits and the flow of information among neurons. For instance, intrinsic time delays within neurons—such as the intervals between spikes or peaks in electrophysiological responses within a single neuron [1]—can vary over time and reveal fluctuations in neuronal activity, contributing to our understanding of how individual neurons respond to stimuli. Another example is the delay in signal transmission during synaptic events between neurons—the time it takes for a presynaptic neuron to trigger a response in a postsynaptic neuron [2]. This signal transmission typically takes milliseconds, may vary depending on synaptic strength or modulation, and is crucial for understanding synaptic efficiency and the synchronization of neural circuits. Similarly, time delays between neural and behavioral responses can also vary over time, as signals travel between the brain and body—such as during motor commands or reflexes, where signals pass through the spinal cord to execute movements.

Biologists have long recognized the biological importance of time delays in neural processes. For instance, owls use auditory time delays to determine target distances during echolocation, where the “best delay” between a pulse and its echo corresponds directly to the prey location [3]. Similarly, time delays play a key role in synfire chains, which are neural circuits that propagate synchronous volleys of spikes, critical for computation in the brain [4]. While these studies have provided insights into how time delays function in groups of neurons, they relied on low-resolution methods that could not capture subtle or time-varying delays at the fine spatiotemporal scales, and often used simplified estimation approaches.

Recent advances in measurement techniques have expanded the capacity to record neural dynamics across scales. High-temporal-resolution electrophysiological recordings, such as depth electrodes [5] and Neuropixels [6], enable the simultaneous recording of thousands of neural activities with sub-millisecond temporal resolution, providing unprecedented insight into time delays at the population neuron levels. In addition, Patch-seq leverages whole-cell patch-clamp electrophysiological recordings with sub-millisecond temporal resolution and integrates these with single-cell RNA sequencing to profile the electrophysiological and transcriptomic properties at single-cell resolution [1]. However, high-sampling-rate electrophysiological and multimodal techniques pose significant analytical challenges for time-varying delay inference due to their high temporal resolution, noise, and multiscale temporal dynamics, making it difficult to extract meaningful timing patterns. Previous studies have proposed multiple approaches to analyze neural timing or align time-series data.

One widely used method is Dynamic Time Warping (DTW), which aligns two time-series by minimizing the global alignment error through dynamic programming [7]. DTW can handle variable-length time-series and is robust to time delays, making it useful in contexts such as speech recognition and activity monitoring. However, it is sensitive to noise, shape, and dynamic differences, and its computational cost is high, though faster approximations exist. Another approach is Canonical Correlation Analysis (CCA), which maximizes the correlation between two sets of signals, reducing dimensionality while preserving meaningful correlations [8]. This makes CCA useful for understanding relationships between multimodal data sources like neural signals and behaviors, but it assumes linear relationships between signals, which limits its effectiveness for nonlinear interactions. The Canonical Time Warping (CTW) method combines DTW and CCA to enable temporal alignment and feature selection for sequences with varying dimensionality [9]. However, it struggles with non-linear transformations and complex temporal dynamics, and its computational burden restricts large-scale applications. Other methods, such as Fourier-based Correlation, leverage the fast Fourier transform (FFT) to align signals based on frequency domain representations [10]. This approach is fast and facilitates noise reduction in the frequency domain, although it assumes linear delays and may struggle with non-periodic signals. Finally, AlignNet, a deep learning-based method, provides end-to-end learning for aligning audio-visual sequences [11]. However, AlignNet is primarily targeted at video and audio alignment in computer vision, using warping and affinity functions that are not applicable for neural time-series data. It is also computationally expensive, tested on low-sampling-rate video and audio, and can be challenging to integrate with domain-specific priors.

To address those challenges, we developed NeuroTD, a sliding-window based alignment approach to analyze time-varying delays in neural activities. By sliding one time series relative to another and computing the mean squared error over each window position, NeuroTD identified the time delay that best aligned the signals, enabling robust, sub-millisecond inference of neural delays even under high noise levels. We evaluated NeuroTD using both simulations and real datasets, including electrophysiological recordings from depth electrodes and Patch-seq data linking gene expression to time-delay features.

## 2 Methods

Our **Fig. 1** provides an overview of the NeuroTD approach for inferring time-varying neural delays. In this conceptual example, two input signals (**x**, red; **y**, blue) contain analogous spike events that occur at different times with varying delays. NeuroTD uses a sliding-window strategy that divides the signals into short overlapping time windows and optimizes their alignment within each window independently.

**Fig. 1.**
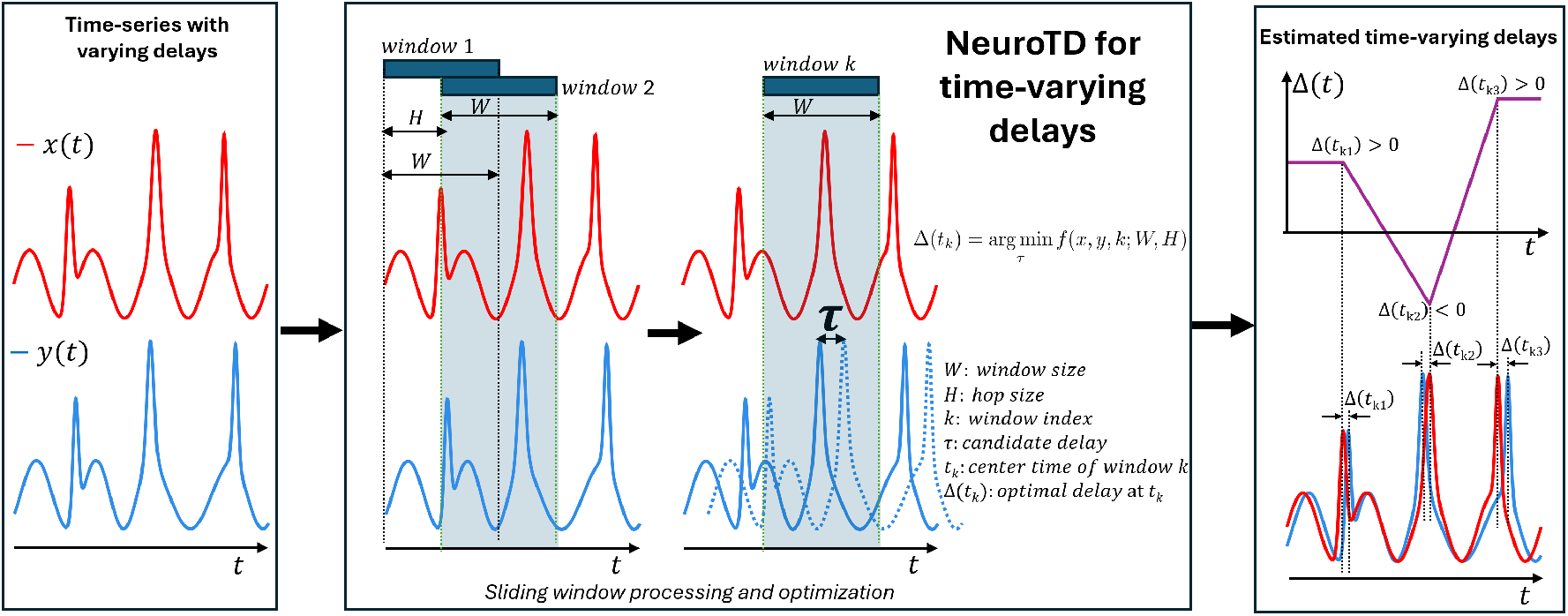
NeuroTD infers time varying delays in neural activities by adaptive sliding window alignment. Input – Time-series with varying delays: Two time-series signals (for example, an electrophysiological recording **x** in red and another electrophysiological or behavioral signal **y** in blue) originate from different sources and may have distinct sampling rates. The signals contain corresponding events that are misaligned in time, potentially due to underlying neural dynamics. In this illustrative example, both **x** and **y** exhibit three analogous spike events (peaks) with varying time offsets, magnitudes, and shapes to be aligned. **NeuroTD: Sliding window processing and optimization:** To capture time-varying delay dynamics, a sliding window of length *W* is applied across both signals. The window moves across time (gray shading) with a hop size *H*, producing overlapping segments of **x** and **y**. For a given window position *k*, the segment of **x** is treated as fixed, while the segment of **y** is shifted by a candidate time delay *τ*. Minimizing the mean squared error between the two windowed segments yields a local delay estimate 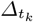 for window *k*. **Output – Estimated time-varying delays with quality score:** Compiling the delay estimates from all windows produces a delay function *Δ*(*t*) (purple curve, top panel) that captures how the relative timing between **x** and **y** evolves over the duration of the signals. The bottom panel illustrates the time delays for the three example spike events: dotted lines indicate event times in **x** and **y**, and horizontal gaps between corresponding dotted lines represent delay magnitudes. Event times on the delay plot *Δ*(*t*) are referenced to the timeline of **x**. Positive delay values (*Δ*_*t*_ *>* 0) indicate that **x** leads **y**, whereas negative values (*Δ*_*t*_ *<* 0) indicate that **x** lags behind.

**Fig. 2.**
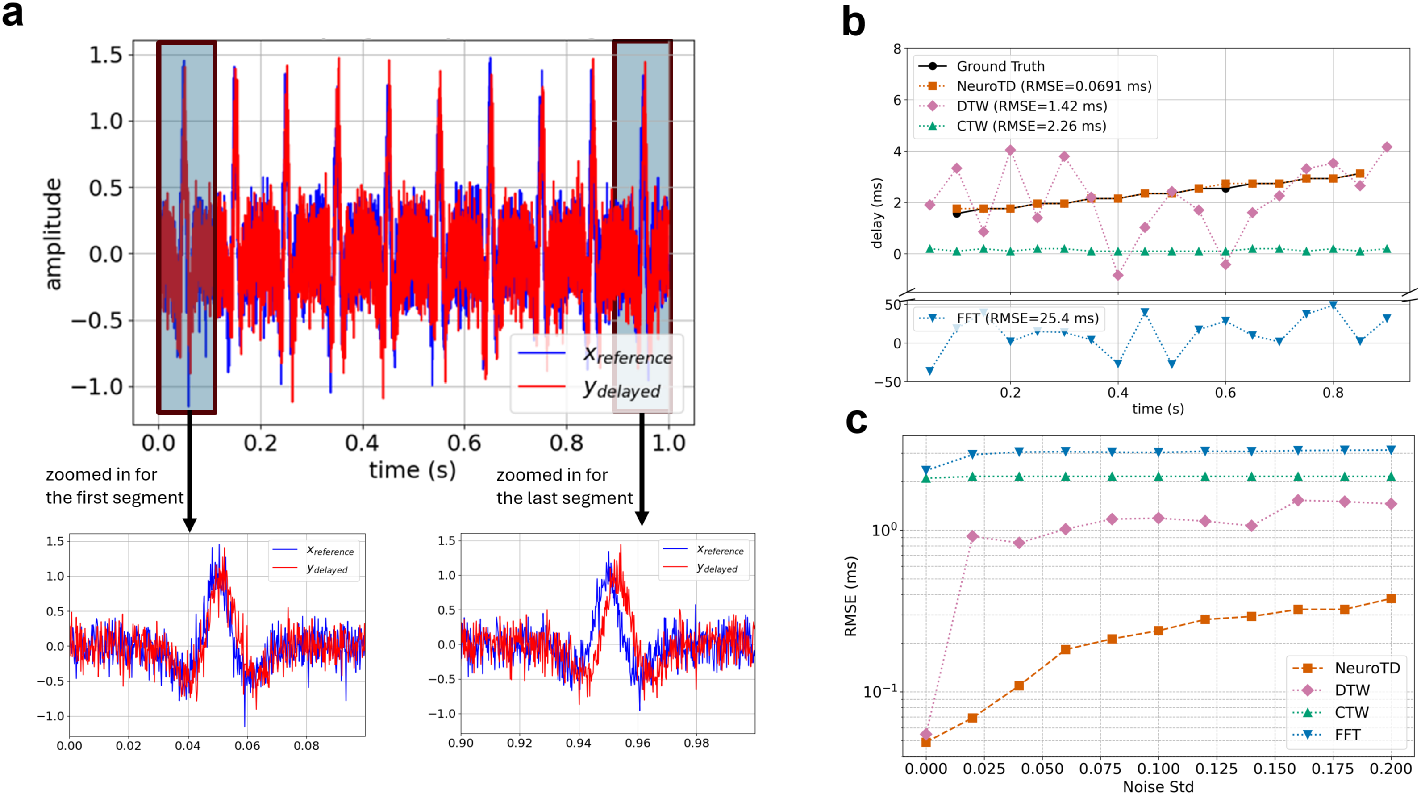
Simulation study on discrete time-varying delays. **a**. Two time-series were simulated according to Section 2.5, *x* (blue, reference) and *y* (red), with piecewise time-varying delays, where *y* is a delayed version of *x*. The zoomed-in sections illustrate delays of 1.56 ms at the beginning and 3.32 ms at the end. Gaussian noise was added at multiple levels to mimic realistic measurement conditions. **b**. Estimated time-varying delays at a fixed noise level (std = 0.020). NeuroTD closely tracked the ground-truth delay profile across time, whereas Dynamic Time Warping (DTW), Canonical Time Warping (CTW), and FFT-based methods exhibited larger deviations. Note: The y-axis is broken to accommodate the large dynamic range of the FFT-based estimates. **c**. NeuroTD delay-estimation error across noise levels with quality-selected window size. For each noise level, the sliding-window size was automatically selected by minimizing the NeuroTD alignment quality score, and the corresponding ground-truth RMSE is reported, demonstrating robust performance without manual tuning across a wide range of noise levels.

For each window, the algorithm identifies the time shift *Δ* that minimizes the mean squared error (MSE) between the segment of *x* and the corresponding segment of *y* shifted by *Δ*. This optimal shift is treated as the local delay estimate for that window. The hop size *H* determines the temporal resolution of the delay estimates, with smaller *H* yielding finer resolution. The window length *W* governs the trade-off between capturing fine-grained temporal dynamics and ensuring robust estimation; in our approach, *W* can be tuned automatically based on the quality score to achieve an optimal balance.

As the window slides across the entire signal, the collection of local delay estimates yields a delay profile *Δ*(*t*) that tracks how the relative timing between *x* and *y* evolves over time. This resulting delay profile provides an efficient and interpretable characterization of neural timing dynamics.

In the following sections, we detail each step of the NeuroTD pipeline.

### 2.1 Input – Time-series with varying delays

NeuroTD takes as input of any two time-series signals, **x** and **y**, which may from electrophysiological recordings or other sources. These signals are assumed to contain corresponding events, but their timing is not strictly aligned—due to neural variability, the delay between corresponding events can change over time. In **Fig. 1**, we illustrate an example where segments of **x** and **y** exhibit three analogous spike events (peaks) with varying time offsets, magnitudes, and shapes that NeuroTD aims to align.

### 2.2 Sliding window processing and optimization

To handle time-varying delays across time-series signals, we first apply a sliding window of length *W* with hop size *H* across both inputs **x** and **y**. Here, *W* is a tunable window length (e.g., *W* = 1023 samples which is approximately 1*/*32 second for 32 kHz-sampled data), and *H* is the hop size, defined as the number of samples between adjacent windows, which determines the stride between successive windows. A common choice is *H* = *W//*2 (where *//* is integer division), but smaller or larger values can be used depending on the desired temporal resolution. For instance, *H* = 1 yields maximal temporal precision (shifts computed at every sample), whereas larger values reduce computational load at the expense of granularity. We restricted *H* ≤ *W/*2 in this paper to maintain adequate resolution.

For each window position, paired local segments of size *W* are extracted from **x** and **y** to align. We denote the windowed signals as *x*[*n*] and *y*[*n*] for simplicity.

Within each window, we align *y* to *x* by minimizing the local mean squared error (MSE) with respect to the shift variable *τ*. Specifically, for the *k*th window with a given starting index *kH*, we define:

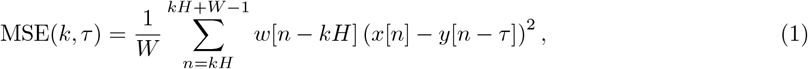

where *w*[*n*] is chosen as a rectangular function (*w*[*n*] = 1) or a Hann function of length *W*:

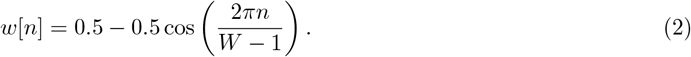

To limit edge effects and enforce locality, the candidate shift range is constrained to |*τ* | ≤ *W/*4. Thus, the optimal delay for the *k*th window is defined as

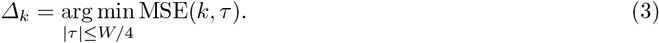

Collecting all *Δ*_*k*_ across windows yields the time-varying delay profile *Δ*(*t*), which captures how the relative timing between **x** and **y** evolves over time.

When *w*[*n*] is chosen as a rectangular window, the MSE can be expanded into three terms to expedite computation:

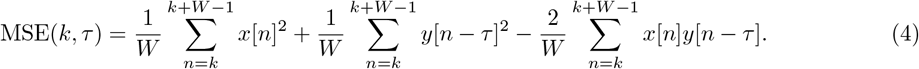

The first two terms, which capture local signal energy, can be precomputed using partial sums. The third term reflects local cross-correlation and can be accelerated via dynamic programming or FFT-based convolution.

#### Tradeoffs

The choice of window size *W* reflects a balance between temporal localization and noise robustness. Larger windows smooth local fluctuations and yield stable estimates, but may blur finer dynamics. Smaller windows enhance spatial resolution but increase sensitivity to noise. Similarly, the hop size *H* controls the temporal density of estimated shifts: smaller *H* improves resolution at the cost of higher computational cost, whereas larger *H* trades off precision for speed. In practice, both *W* and *H* are selected based on expected delay variability and signal quality.

### 2.3 Output – Time-varying delays

The output of NeuroTD is a time-resolved sequence of estimated delays *Δ*(*t*_*k*_) for *k* = 1 · · · *K*. Each *Δ*(*t*_*k*_) represents the relative delay between signals **x** and **y** at window *k*. By convention, a positive delay (*Δ*(*t*_*k*_) *>* 0) indicates that **x** leads **y** at time *t*_*k*_, whereas a negative delay (*Δ*(*t*_*k*_) *<* 0) indicates that **x** lags behind. This dynamic delay profile, *Δ*(*t*), captures how the timing relationship between the two signals evolves over time, providing insight into neural signal propagation and coordination. For example, fluctuations in *Δ*(*t*) can be related to behavioral or physiological event, or compared against ground-truth timings, when available. In downstream analysis, the delay sequence can also be post-processed (e.g., smoothed or thresholded, or weighted by a confidence measure) depending on the noise level and the specific application.

### 2.4 Evaluation and quality score

To evaluate the performance of NeuroTD, we use the root-mean-square error (RMSE) as the primary metric for quantifying the difference between the predicted time delay *Δt*\* and the ground truth *Δt*. It is computed as 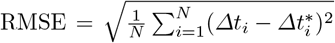, where *N* is the number of samples. A lower RMSE indicates better alignment between the time-series signals and more accurate time-delay estimation.

In addition, we report an *alignment quality* score *q*, which measures the consistency of alignment after applying *Δt*\*. For each sliding window *m*, we first compute a normalized root-mean-square error (NRMSE) between *x* and the shifted *y*:

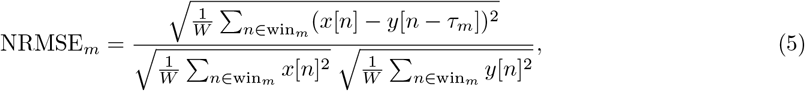

where *W* is the window length and *τ*_*m*_ is the estimated shift for window *m*. The overall alignment quality is then defined as

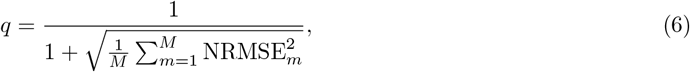

where *M* is the number of valid windows. Higher *q* indicates better alignment consistency and is used to select the optimal window size that maximizes quality. The score ranges from 0 *< q* ≤ 1, where *q* = 1 represents perfect alignment between the signals.

To determine the optimal window size, we compute the alignment quality *q* for a range of candidate odd window sizes (e.g., 31, 63, 127, 255 samples), ensuring that each window has an integer center point. For each size, NeuroTD performs sliding-window alignment and outputs the corresponding *q*. We then select the window size that maximizes *q*, denoted as *W* * = arg max_*W*_ *q*(*W*). This procedure ensures that the chosen window balances temporal resolution and estimation stability: smaller windows capture rapid delay changes but may increase noise sensitivity, whereas larger windows improve robustness but may smooth over short-term dynamics. The quality-based selection therefore provides a principled, data-driven criterion for adapting the window length to the characteristics of the analyzed signals.

### 2.5 Simulation design

To evaluate NeuroTD across a broad range of biologically relevant timing regimes, we designed two complementary simulation settings. The discrete-delay simulation models sparse or event-driven processes with abrupt delay changes, whereas the continuous-delay simulation captures smoothly evolving delays characteristic of ongoing neural dynamics; together, these simulations assess robustness under both piecewise and smoothly varying temporal structures.

#### Discrete delay simulation

We constructed a pair of signals (*x*(*t*), *y*(*t*)) on a normalized time interval *t* ∈ [0, 1) to validate time-delay estimation. The reference signal *x*(*t*) was designed with structured features, e.g. a sinusoidal waveform superimposed with a spike-like transient, to provide both oscillatory and abrupt signal components. The signal *y*(*t*) was then generated as a piecewise-delayed version of *x*(*t*): the time interval was partitioned into *N*_seg_ equal segments, and in each segment *i, y*(*t*) was a time-shifted copy of *x*(*t*) with a fixed delay *Δ*_*i*_. The delays *Δ*_*i*_ were chosen to increase monotonically with *i*, approximating a linearly increasing delay trend over time. Let *Δ*_min_ and *Δ*_max_ denote the delays in the first and last segments respectively, the intermediate delays were assigned via linear interpolation:

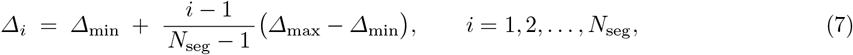

ensuring that *Δ*_1_ = *Δ*_min_ and 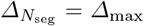. Within each segment, the delay was held constant at *Δ*_*i*_, so that *y*(*t*) = *x*(*t* − *Δ*_*i*_) for all *t* in segment *i*, following the same sign convention as defined in Section 2.3 (positive: *x* leads *y*).

In this paper, we used *N*_seg_ = 10 segments, with delays monotonically increasing from approximately *Δ*_min_ = 1.56 ms at *t* = 0 to *Δ*_max_ = 3.32 ms at *t* = 1(*s*), yielding a piecewise-linear delay profile across the signal duration. To simulate realistic conditions, independent Gaussian noise was added to both *x*(*t*) and *y*(*t*). We evaluated NeuroTD against multiple baseline methods, including Dynamic Time Warping (DTW), Canonical Time Warping (CTW), and an FFT-based cross-correlation approach, under varying noise levels. Each method attempted to recover the known time-delay function *Δ*(*t*), and accuracy was quantified using the root-mean-square error (RMSE) between the estimated and true delay trajectories. In our experiments, the noise standard deviation was varied across a range from 0 to 0.20 in evenly spaced steps.

#### Continuous delay simulation

We next simulated a pair of signals (*x*(*t*), *y*(*t*)) over *t* ∈[0, 1) where *y* was a smoothly delayed version of *x* via a time-varying delay curve *Δ*(*t*). The reference *x*(*t*) was composed of a low-amplitude sinusoid and *N*_spike_ Gaussian spike transients to capture both ongoing rhythmic activity and transient event structure commonly observed in neural responses to stimuli:

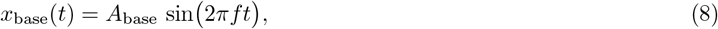

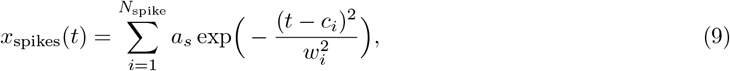

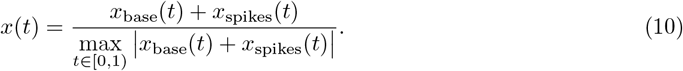

Where *A*_base_ is the amplitude of the base sinusoid, *f* is its frequency, *a*_*s*_ is the spike amplitude, *c*_*i*_ is the spike center, *w*_*i*_ controls the width of spike *i*, and the maximum of *x*(*t*) is normalized to 1.

To generate the delayed signal, we prescribed a smooth delay curve

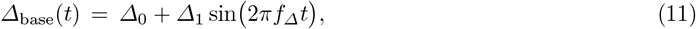

and optionally perturbed it at spike centers to mimic event-specific delays. Let *ε*_*i*_ denote small per-spike perturbations (e.g., i.i.d. zero-mean) at spike centers 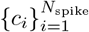. We then fit a cubic spline *Δ*(*t*) through the constraints

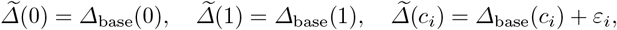

yielding a *C*^2^ delay curve with boundary stability and smooth transitions between edited events. Finally, we warped time with wrap-around

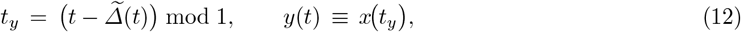

where the modulo operation enforces circular boundaries so that samples exceeding *t* = 1 wrap to *t* = 0, and the delay sign follows the convention in Section 2.3.

Unless otherwise specified, spike centers *c*_*i*_ are roughly uniformly spaced on [0, 1) with small jitter, and spike widths *w*_*i*_ are varied mildly to vary morphology. This continuous-delay setup complements the discrete-segment simulation (Section 3.1) by testing algorithms under smoothly varying, event-perturbed lags, while preserving known ground truth at all *t* and at the spike times {*c*_*i*_} of salient events.

### 2.6 Implementation and hyperparameters

All simulations and analyses were implemented in Python 3.12 using the NumPy (v2.3.3) and SciPy (v1.16.2) libraries for signal generation and numerical computations. All computations were performed on a standard desktop equipped with an Intel i9-11900 CPU and 64 GB RAM.

NeuroTD estimated time-varying shifts using a sliding-window strategy with window length *W* and hop size *H* as its key hyperparameters. The window size *W* was automatically selected using NeuroTD’s internal quality score, which measures post-alignment consistency. The hop size was set to *H* = ⌊*W/*2⌋ for discrete-delay simulations and real data to ensure computational efficiency, and to *H* = 1 sample for continuous-delay simulations to preserve fine temporal precision.

Each simulation was repeated across multiple noise levels to compute average root-mean-square error (RMSE) and quality scores. The parameter choices for signal generation (e.g., *N*_seg_, *Δ*_min_–*Δ*_max_, *N*_spike_, *A*_base_, *a*_*s*_, *w*_*i*_) are detailed in their respective subsections for transparency.

### 2.7 Real datasets and data processing

We used four datasets - two simulated and two real-world. The simulated datasets included one with discrete, piecewise time-varying delays and another with smoothly varying continuous delays, both designed to benchmark NeuroTD against existing methods under controlled conditions. The real-world datasets spanned different biological and technical scales: (1) a neural circuits dataset that includes neuron electrical activities recorded from 32–64 intracranial recording sites across medial temporal lobe regions, measured using depth electrodes, with their spatial positions recorded to map time delays and signal transmission pathways [5]; and (2) a Patch-seq dataset from mouse motor cortex that combines electrophysiological recordings with single-cell gene expression profiles [12]. We used all valid intratelencephalic (IT) subtype cells with available intrinsic spiking time delay (ISTD) values (237 of 254 total IT cells). Gene expression data were first restricted to protein-coding genes. Raw count matrices were normalized by gene length to account for differences in transcript size, and normalized values were transformed using log_2_(value + 1) to stabilize variance across expression levels. To focus analyses on informative transcriptional features, genes with low expression prevalence were filtered out, retaining 7,988 genes that were expressed in at least 50 % of cells from the original set of 18,668 protein-coding genes.

## 3 Results

### 3.1 Simulation study on discrete time-varying delays

In this simulation study, we generated two synthetic signals with dynamically changing time delays to evaluate the effectiveness of NeuroTD. **Section 3.1a** illustrated the simulated signals, with delays increasing from 1.56 ms at the beginning to 3.32 ms at the end of the sequence. We simulated 10 different delays for each of the segments. To replicate realistic recording conditions, Gaussian noise was added at eleven levels, with standard deviations linearly spaced between 0 and 0.20. For each noise level, NeuroTD estimated time delays and alignment quality scores using six candidate window sizes {33, 65, 129, 257, 513, 1025} samples, following an odd-length sequence of the form 2^*n*^ + 1. These window sizes correspond to durations ranging from 6.45 to 200.20 ms. The window size yielding the highest quality score was selected as the final result.

The results in **Section 3.1b** and **Section 3.1c** showed that NeuroTD consistently achieves accurate alignment and time-delay estimation, even under high noise. At a fixed noise level (std = 0.020) in **Section 3.1b**, NeuroTD achieved an RMSE of 0.0691 ms, closely tracking the ground-truth delay profile. In contrast, Dynamic Time Warping (DTW) yielded an RMSE of 1.42 ms, Canonical Time Warping (CTW) 2.26 ms, and the FFT-based method 25.4 ms. Across increasing noise levels (**Section 3.1c**), NeuroTD consistently outperformed all baseline methods, maintaining low delay-estimation error through automatic selection of the sliding-window size based on a data-driven quality score, *without access to ground-truth delay information*. For DTW, CTW, and FFT, the reported RMSE at each noise level corresponds to the minimum value obtained after sweeping all candidate window sizes *with access to ground-truth delay information*, representing an *oracle selection most favorable to the baseline methods*. In contrast, NeuroTD reports RMSE from the configuration selected solely by minimizing the alignment quality criterion. Additional benchmarking results, including NeuroTD quality scores, evaluated and selected window sizes at each noise level, RMSE values across all window sizes, and oracle versus quality-selected RMSE comparisons across methods, are provided in **Supplementary Tables S1**.

### 3.2 Simulation study of continuous time-varying delays

In this continuous-delay simulation, we evaluated NeuroTD’s ability to track smoothly varying time shifts. Following the design in Section 2.5, we set *N*_spike_ = 10, *f* = 4, *Δ*_0_ = 0.00, *Δ*_1_ = 0.002, *f*_*Δ*_ = 0.1, *A*_base_ = 0.1, and *a*_*s*_ = 0.5. The spike centers *c*_*i*_ were approximately uniformly spaced over [0, 1) with Gaussian jitter of standard deviation 0.015, and spike widths were drawn from the range *w*_*i*_ ∈ (0.005, 0.01) seconds. At each spike center, an independent Gaussian perturbation was applied to the delay value with standard deviation 0.001 s, equivalent to 1 ms. Gaussian noise was added independently to both *x* and *y* at eleven levels, with standard deviations linearly spaced between 0 and 0.20, to mimic realistic recording conditions. For each noise level, NeuroTD estimated time delays and alignment quality scores using six candidate window sizes {33, 65, 129, 257, 513, 1025} samples, following an odd-length sequence of the form 2^*n*^ +1. These window sizes correspond to durations ranging from 6.45 to 200.20 ms. The window size yielding the highest quality score was selected as the final result. The hop size was fixed to one sample to ensure high temporal resolution in time-delay estimation.

As shown in **Fig. 3a**, the true delay curve increased gradually across the signal, with realistic Gaussian spikes and a sinusoidal baseline to mimic electrophysiological recordings. Results in **Fig. 3b** show the estimated delay trajectories at a fixed noise level (std = 0.02). NeuroTD was evaluated using quality-selected window sizes without access to ground-truth delays, whereas DTW, CTW, and the FFT-based method were evaluated under oracle window-size selection with access to ground-truth delays. Under this setting, NeuroTD reliably recovered the smoothly varying continuous delay profile and closely followed the ground truth across time (RMSE = 0.29 ms), while DTW (RMSE = 0.69 ms) and CTW (RMSE = 1.32 ms) exhibited substantially larger deviations and the FFT-based approach produced highly unstable estimates with very large error (RMSE = 11.61 ms), demonstrating that NeuroTD effectively adapts to gradual continuous delay variations whereas baseline methods fail to capture fine-grained temporal dynamics. To evaluate robustness under smoothly varying delays, we analyzed RMSE as a function of noise level in the continuous-delay simulation (**Fig. 3c**). For each of eleven noise levels, DTW, CTW, FFT, and NeuroTD were first evaluated under oracle window-size selection, where the window minimizing RMSE was chosen with access to ground-truth delays. Under this oracle setting, NeuroTD achieved the lowest RMSE across all noise levels, providing an upper bound on its achievable performance. We next evaluated NeuroTD under a deliberately unfavorable setting in which the window size was selected solely by maximizing the internal alignment quality score, without access to ground-truth information. Under this quality-based selection, NeuroTD continued to outperform DTW, CTW, and FFT for all nonzero noise levels. A small performance gap was observed only at zero noise, where the window size that minimizes RMSE differs from the window size selected by the alignment quality criterion. Additional benchmarking results, including NeuroTD quality scores, evaluated and selected window sizes at each noise level, RMSE values across all window sizes, and oracle versus quality-selected RMSE comparisons across methods, are provided in **Supplementary Tables S2**.

**Fig. 3.**
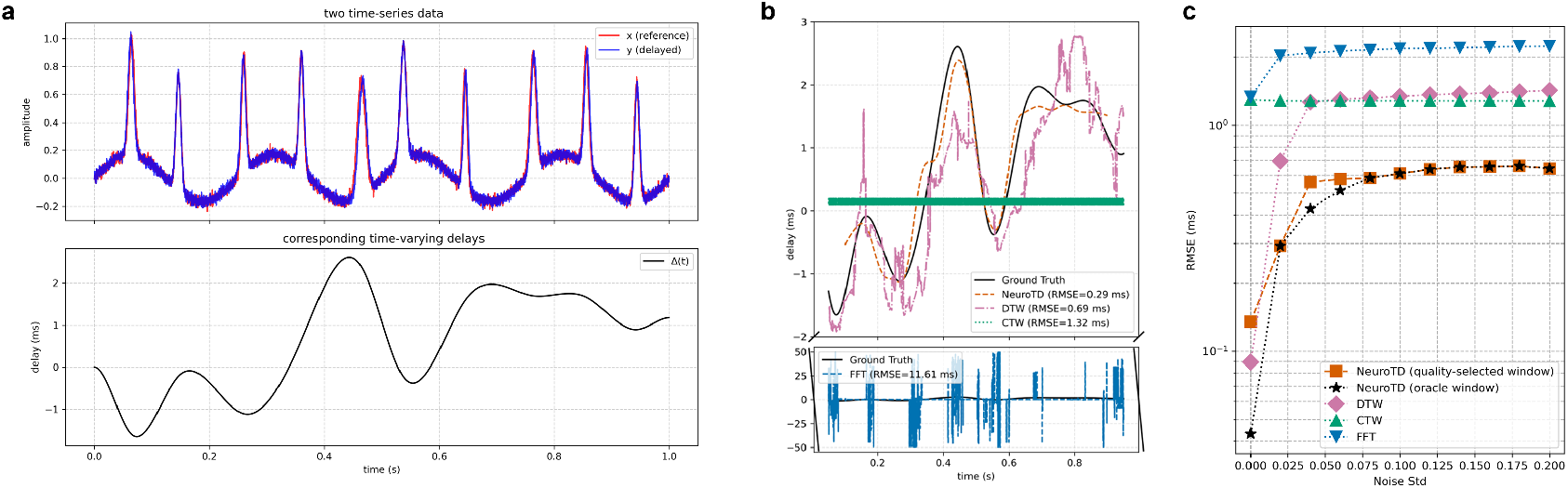
Simulation study of continuous time-varying delays. **a**. A smoothly varying time delay *Δ*(*t*) was imposed between two signals *x*(*t*) and *y*(*t*), generated by combining a sinusoidal baseline with *N*_spike_ = 10 Gaussian spikes. The delay increased continuously from 0.0 ms to 10.0 ms across the signal duration. **b**. Estimated time-varying delays at a fixed noise level (std = 0.02). NeuroTD closely tracked the ground-truth continuous delay trajectory across time, maintaining sub-millisecond accuracy. In contrast, Dynamic Time Warping (DTW) and Canonical Time Warping (CTW) exhibited larger deviations, while the FFT-based method showed unstable estimates with large dynamic range. Note: The y-axis is broken to accommodate the large dynamic range of the FFT-based estimates. **c**. Root-mean-square error (RMSE) as a function of noise level for the continuous-delay simulation. DTW, CTW, FFT, and NeuroTD (oracle window) report the minimum RMSE across candidate window sizes using access to ground-truth delays, whereas NeuroTD (quality-selected window) selects window size using only the internal alignment quality score, without access to ground-truth delays.

**Fig. 4.**
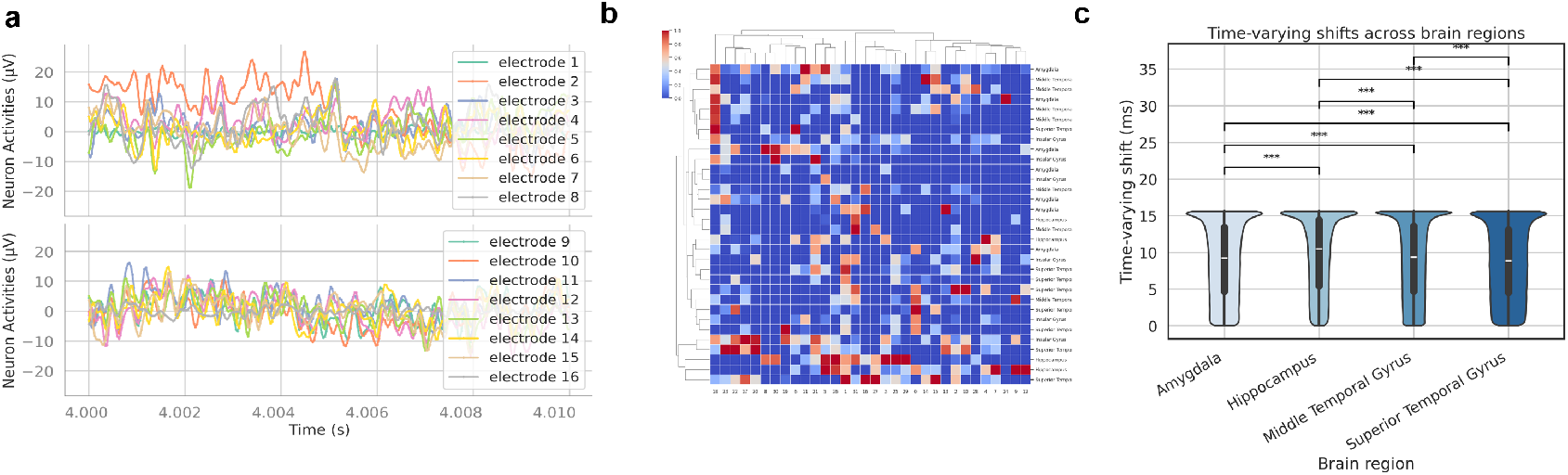
Time delays of neural electrical activities associated with 3D positions across the human medial temporal lobe. **a**. Simultaneously recorded neural electrical activities from 16 depth electrodes. A 10 ms segment of neural activity recorded from 16 of 32 depth electrodes across medial temporal lobe regions [5], illustrating variability in iEEG signal timing and waveform shape. **b**. Pairwise time-delay matrix among 32 iEEG channels (electrode contacts). A heatmap of estimated time delays between all channel pairs, with hierarchical clustering highlighting temporal similarity across channels. **c**. Distribution of time-varying delays across brain regions. Violin plots show delay distributions stratified by anatomical region. Delays in the hippocampus were significantly larger than those in other sampled medial temporal lobe regions (Bonferroni-corrected two-sided Welch’s t-tests on per-delay values, *p <* 0.001).

### 3.3 Time delays of neural electrical activities associated with 3D positions across the human medial temporal lobe

We applied NeuroTD to intracranial electroencephalography (iEEG) recorded using 32 to 64 depth electrodes across medial temporal lobe regions [5]. These data were acquired while human subjects performed a modified Sternberg verbal working memory task — memorizing a set of letters during an encoding period, maintaining them over a delay, and responding to a probe letter — allowing analysis of brain activity during working memory maintenance. Each iEEG channel was recorded simultaneously and assigned a 3D anatomical position, allowing for spatiotemporal analysis of signal transmission. **Section 3.3a** showed a 10 ms segment of neural activity from 16 electrode contacts, illustrating variability in iEEG signal timing and waveform shape. NeuroTD was then used to estimate pairwise time delays between all iEEG channels (depth-electrode contacts), summarized as a clustered delay matrix in **Section 3.3b**. In **Section 3.3c**, we grouped the delays by brain region, pooling data across all experimental sessions. The hippocampus exhibited significantly larger time-varying delays than other sampled medial temporal lobe regions (Bonferroni-corrected two-sided Welch’s t-tests on per-delay values, *p <* 0.001). This may reflect slower local dynamics or the layered microcircuit structure of the hippocampus. However, prior studies have reported synchronized activity between the hippocampus and prefrontal cortex during working memory tasks, without necessarily indicating slow signal transmission [13]. Similarly, feedforward and feedback pathways in cortical circuits have been shown to use distinct frequency channels, but not slower conduction [14]. The observed ranking also showed the superior temporal gyrus had the lowest delays, while the amygdala and middle temporal gyrus had delays between hippocampus and superior temporal gyrus. These region-specific delay rankings and statistical differences were consistent across individual experimental sessions, as shown in **Supplementary Figure S1**. Together, these results demonstrated that NeuroTD can extract fine-grained temporal information from deep-brain activity and revealed region-specific timing patterns, while emphasizing the importance of cautious interpretation.

### 3.4 Gene expression correlations with electrophysiological time-delays of excitatory neurons in mouse motor cortex

In this application, we examined the relationship between time delays in electrophysiological responses and gene expression in the mouse motor cortex using data of excitatory neurons from Scala et al. [12]. **Fig. 5a** illustrated the inter-spike intervals computed by NeuroTD, or the time gaps between consecutive spikes, recorded during neural activity sweeps. These gaps changed in response to neural stimuli and indicated how neurons adjusted their firing patterns. **Fig. 5b** quantified these changes by computing the slope of the ISI changes over multiple sweeps. The slopes reflected how quickly or slowly the time delays evolved, offering insights into the dynamics of neuronal firing patterns. The inter-spike time delays (ISTD) for each neuron was computed as the mean slope across sweeps, capturing robust neural dynamics. As illustrated in **Fig. 5c**, we correlated ISTDs with gene expression levels of 7,988 highly expressed genes across all 237 excitatory neurons (details in Section 2.7). **Fig. 5d** revealed the top 20 genes with the strongest absolute correlations (both positive and negative) with the ISTD time-delay features for those IT cells. This correlation helped identify genes potentially associated with rapid changes in neural response, providing a genetic link to neural firing dynamics. We performed an enrichment analysis of the top 200 correlated genes ranked by the absolute value of their correlation with ISTD scores (**Fig. 5e**), which highlighted several biological processes relevant to electrophysiological signaling. These include: GO:0050804 (modulation of chemical synaptic transmission, *p*-value 10^−6.2^), reflecting direct regulation of synaptic communication; R-MMU-112316 (the Neuronal System Reactome pathway, *p*-value 10^−4.6^), encompassing core neuronal signaling processes; GO:0051480 (regulation of cytosolic calcium ion concentration, *p*-value 10^−4.0^), highlighting calcium-dependent control of neuronal excitability; GO:0048812 (neuron projection morphogenesis, *p*-value 10^−3.6^), implicating structural organization of neuronal projections relevant to signal propagation; and GO:0007616 (long-term memory, *p*-value 10^−3.2^), linking electrophysiological variability to higher-order neural function. Together, these enriched categories emphasize synaptic transmission, ionic regulation, and neuronal signaling mechanisms underlying electrophysiological time-delay dynamics. The full list of the top 200 genes ranked by the absolute value of their correlation with ISTD scores, including both positive and negative associations, is provided as **Supplementary Data S1**.

**Fig. 5.**
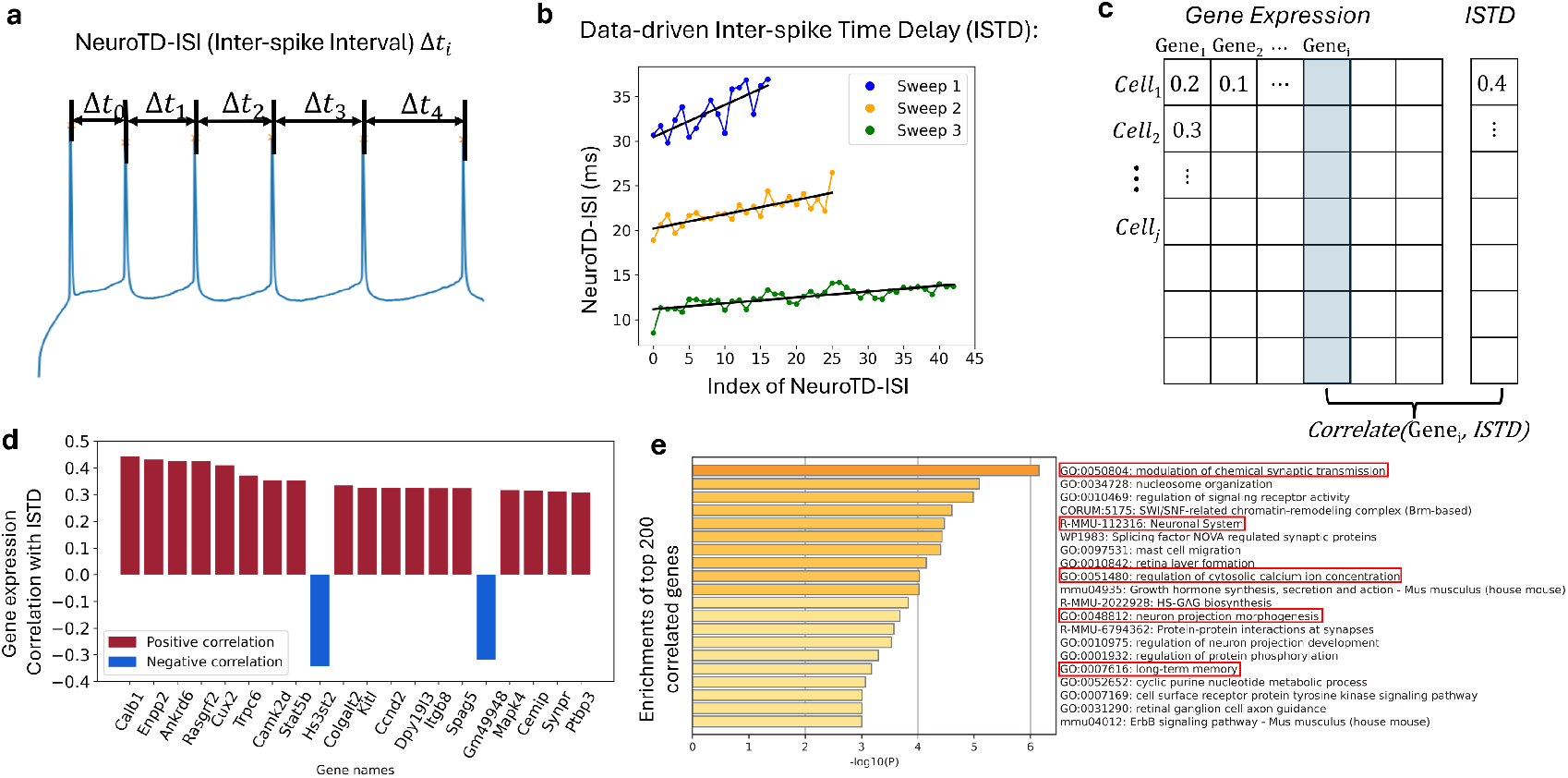
Gene expression correlations with electrophysiological time-delay features in mouse motor cortex. **a**. NeuroTD computed inter-spike intervals. A representative electrophysiological sweep illustrated time gaps between consecutive spikes (*Δt*_*i*_) computed using NeuroTD. **b**. Data-driven inter-spike time delays (ISTDs). Changes in inter-spike intervals across multiple sweeps were quantified by computing the mean slope of ISI progression for each neuron, yielding the ISTD values. **c**. Gene expression and time-delay features. A matrix representation of single-cell gene expression profiles aligned with corresponding ISTDs for each neuron. **d**. Correlation between gene expression and ISTDs. Bar plots showed the top 20 genes most strongly correlated with ISTD values, with red indicating positive correlations and blue indicating negative correlations. **e**. Functional enrichment of ISTD-associated genes. Enrichment analysis of the top 200 genes ranked by the absolute value of their correlation with ISTDs highlighted Gene Ontology and pathway terms related to synaptic transmission, neuronal signaling, and excitatory postsynaptic regulation.

## 4 Discussion

In this work, we introduced NeuroTD, a sliding-window alignment approach for analyzing time-varying delays in neural electrical activities. NeuroTD combines local alignment with adaptive window-size tuning to capture dynamic sub-millisecond delays while remaining robust to noise, and provides confidence scoring to quantify the reliability of inferred delays. In real datasets, NeuroTD estimated time delays in intracranial electroencephalography (iEEG) recordings from the human hippocampus, revealing region-specific delay patterns among neural signals, where future graph-based analyses of delay structures may provide novel insights into circuit-level interactions. We also applied NeuroTD to Patch-seq data, linking time delays in electrophysiological recordings with gene expression, and found that delay-associated genes were enriched for pathways related to ion transport and neuronal excitability, indicating that these processes are associated with variability in neuronal signaling.

NeuroTD assumes that the analyzed signals share common events to align; extreme cases with entirely non-overlapping events or significant recording artifacts could challenge the method. In practice, windows with insufficient signal content typically yield low alignment quality scores and should be down-weighted or excluded from downstream analysis.

Although NeuroTD offers high temporal resolution for noisy signals, certain limitations remain. In the current implementation, window-level delays are estimated by minimizing a local error criterion and can optionally be smoothed into a continuous delay curve using overlap-weighted reconstruction across adjacent windows. This reduces visible jitter but does not fully eliminate noise-driven variability in the underlying window-wise delay estimates. Future extensions could incorporate explicit temporal continuity constraints or probabilistic formulations to further stabilize delay trajectories, while balancing improved robustness against additional computational cost.

In the hippocampal recordings of Section 3.3, depth electrodes had linearly arranged, evenly spaced contacts, providing limited distance variation within each region. Accordingly, delays examined as a function of contact spacing showed no clear trend. Future studies using denser or more uniformly distributed electrodes will be required to clarify how anatomical distance contributes to signal-propagation delays. The observed region-specific delay rankings should therefore be interpreted cautiously, as they may reflect differences in firing statistics, network states, or sample coverage rather than intrinsic differences in signal conduction.

NeuroTD processed the full Patch-seq dataset Section 3.4(11.1 GB compressed, 36 GB in-memory) within minutes on a standard desktop machine, illustrating practical runtime efficiency. Since computational cost scales with signal length and the number of sliding windows, future improvements such as pre-filtering inactive periods could further enhance scalability for long or sparsely active recordings.

## 5 Data Availability

Simulated data is publicly available on GitHub at https://github.com/daifengwanglab/NeuroTD. Electrophysiological recordings from depth electrodes in the human medial temporal lobe are provided by [5] and available at https://dandiarchive.org/dandiset/000574. Electrophysiological recordings and gene expression data from mouse motor cortex neurons, collected using Patch-seq, are provided by [12] and available at https://dandiarchive.org/dandiset/000008.

## 6 Code Availability

The NeuroTD code is implemented in Python, and the complete NeuroTD package, along with example implementations, are publicly accessible at https://github.com/daifengwanglab/NeuroTD.

## 7 Acknowledgments and Funding

This work is supported by National Institutes of Health grants, RF1MH128695, R01AG067025, P50HD105353, and National Science Foundation Career Award 2144475.

## 8 Contributions

Study conception, D.W.; Conceptualization, X.H. and D.W.; Methodology, X.H. and D.W.; Formal analysis, X.H.; Investigation, X.H., Q.D., Q.C. D.W., and Q.D.; Data curation, X.H., N.C.K., S.A., and A.Z.L.; Software, X.H.; Writing–Original Draft, X.H.; Writing–Review and Editing, X.H., N.C.K., S.A., Q.D., Q.C., and D.W.; Supervision, D.W.; Funding acquisition, D.W.

