## Supplementary Materials for "NeuroTD infers time varying delays in neural activities by adaptive sliding window alignment"

### Supplementary Information

### Supplementary Table

**Supplementary Table S1: Discrete simulation benchmarking across window sizes and noise levels.** For each noise level, multiple candidate window sizes are evaluated, following an odd-length sequence of the form $2^n+1$, ranging from 33 to 1025 samples (6.45 ms to 200.20 ms). The NeuroTD quality score is used to select the optimal window; the minimum quality score is highlighted in bold, and the corresponding window size and NeuroTD RMSE (alignment error, in ms) are also shown in bold. For each method (NeuroTD, DTW, CTW, and FFT), the minimum RMSE across window sizes within a given noise level is underlined, indicating oracle performance. For all non-zero noise levels, the window selected by the NeuroTD quality score yields the same RMSE as the RMSE-optimal window, demonstrating that quality-based selection recovers the oracle solution without access to ground-truth delay information. At zero noise, the quality-selected RMSE (0.049 ms) differs slightly from the optimal RMSE (0.047 ms), indicating near-optimal but not exact agreement.

| Noise | Window (samples, ms) | Quality | NeuroTD | DTW | CTW | FFT |
|---|---|---|---|---|---|---|
| 0.000 | 33 (6.4) | 201.495 | 0.067 | 0.785 | 2.401 | 2.473 |
| 0.000 | 65 (12.7) | 151.715 | 0.047 | 0.464 | 2.392 | 2.451 |
| 0.000 | 129 (25.2) | 330.352 | 0.067 | 0.249 | 2.373 | 2.428 |
| 0.000 | 257 (50.2) | 4.356 | 0.098 | 0.120 | 2.338 | 2.343 |
| 0.000 | **513 (100.2)** | **0.245** | **0.049** | 0.055 | 2.255 | 36.933 |
| 0.000 | 1025 (200.2) | 0.348 | 81.809 | 0.070 | 2.099 | 100.000 |
| 0.02 | 33 (6.4) | 5.662 | 2.183 | 1.905 | 2.368 | 2.935 |
| 0.02 | 65 (12.7) | 5.418 | 2.830 | 1.933 | 2.363 | 3.707 |
| 0.02 | 129 (25.2) | 4.850 | 4.402 | 1.926 | 2.343 | 4.868 |
| 0.02 | 257 (50.2) | 4.088 | 1.405 | 1.888 | 2.308 | 8.907 |
| 0.02 | **513 (100.2)** | **0.588** | **0.069** | 1.422 | 2.260 | 25.362 |
| 0.02 | 1025 (200.2) | 0.595 | 57.791 | 0.918 | 2.153 | 64.297 |
| 0.04 | 33 (6.4) | 4.132 | 2.262 | 1.969 | 2.366 | 3.046 |
| 0.04 | 65 (12.7) | 4.040 | 2.962 | 2.007 | 2.359 | 3.846 |
| 0.04 | 129 (25.2) | 3.519 | 4.551 | 2.000 | 2.346 | 5.193 |
| 0.04 | 257 (50.2) | 2.948 | 1.497 | 2.059 | 2.312 | 9.033 |
| 0.04 | **513 (100.2)** | **0.815** | **0.109** | 1.293 | 2.260 | 25.811 |
| 0.04 | 1025 (200.2) | 0.818 | 57.791 | 0.835 | 2.153 | 51.494 |
| 0.06 | 33 (6.4) | 3.448 | 2.331 | 2.012 | 2.364 | 3.064 |
| 0.06 | 65 (12.7) | 3.389 | 3.072 | 2.058 | 2.356 | 4.021 |
| 0.06 | 129 (25.2) | 2.968 | 4.556 | 2.087 | 2.342 | 5.269 |
| 0.06 | 257 (50.2) | 2.474 | 1.711 | 2.115 | 2.316 | 8.728 |
| 0.06 | **513 (100.2)** | **0.981** | **0.183** | 1.731 | 2.266 | 26.951 |
| 0.06 | 1025 (200.2) | 0.985 | 57.791 | 1.016 | 2.153 | 58.737 |
| 0.08 | 33 (6.4) | 3.044 | 2.392 | 2.051 | 2.365 | 3.041 |
| 0.08 | 65 (12.7) | 2.988 | 3.131 | 2.068 | 2.356 | 4.135 |
| 0.08 | 129 (25.2) | 2.663 | 4.575 | 2.106 | 2.344 | 5.886 |
| 0.08 | 257 (50.2) | 2.218 | 1.714 | 2.125 | 2.325 | 8.011 |
| 0.08 | **513 (100.2)** | **1.112** | **0.213** | 1.791 | 2.266 | 24.776 |
| 0.08 | 1025 (200.2) | 1.117 | 57.791 | 1.173 | 2.153 | 64.396 |
| 0.1 | 33 (6.4) | 2.765 | 2.407 | 2.091 | 2.367 | 3.030 |
| 0.1 | 65 (12.7) | 2.718 | 3.110 | 2.103 | 2.359 | 4.233 |
| 0.1 | 129 (25.2) | 2.463 | 4.589 | 2.071 | 2.345 | 6.146 |
| 0.1 | 257 (50.2) | 2.061 | 2.362 | 2.075 | 2.325 | 9.321 |
| 0.1 | **513 (100.2)** | **1.216** | **0.239** | 2.076 | 2.266 | 23.895 |
| 0.1 | 1025 (200.2) | 1.223 | 57.791 | 1.187 | 2.153 | 65.268 |
| 0.12 | 33 (6.4) | 2.560 | 2.426 | 2.137 | 2.367 | 3.080 |
| 0.12 | 65 (12.7) | 2.520 | 3.117 | 2.152 | 2.359 | 4.444 |
| 0.12 | 129 (25.2) | 2.319 | 4.593 | 2.140 | 2.344 | 6.412 |
| 0.12 | 257 (50.2) | 1.955 | 2.741 | 2.150 | 2.321 | 10.868 |
| 0.12 | **513 (100.2)** | **1.300** | **0.280** | 2.115 | 2.266 | 23.895 |
| 0.12 | 1025 (200.2) | 1.307 | 57.791 | 1.139 | 2.153 | 64.827 |
| 0.14 | 33 (6.4) | 2.397 | 2.464 | 2.164 | 2.366 | 3.068 |
| 0.14 | 65 (12.7) | 2.369 | 3.169 | 2.185 | 2.358 | 4.430 |
| 0.14 | 129 (25.2) | 2.213 | 4.959 | 2.183 | 2.343 | 6.544 |
| 0.14 | 257 (50.2) | 1.888 | 2.676 | 2.217 | 2.321 | 11.715 |
| 0.14 | **513 (100.2)** | **1.366** | **0.293** | 2.174 | 2.266 | 23.895 |
| 0.14 | 1025 (200.2) | 1.374 | 57.791 | 1.065 | 2.153 | 55.844 |
| 0.16 | 33 (6.4) | 2.269 | 2.478 | 2.188 | 2.365 | 3.113 |
| 0.16 | 65 (12.7) | 2.245 | 3.249 | 2.230 | 2.359 | 4.385 |
| 0.16 | 129 (25.2) | 2.118 | 5.038 | 2.270 | 2.345 | 6.628 |
| 0.16 | 257 (50.2) | 1.828 | 3.185 | 2.296 | 2.321 | 12.211 |
| 0.16 | **513 (100.2)** | **1.419** | **0.324** | 2.401 | 2.266 | 23.840 |
| 0.16 | 1025 (200.2) | 1.430 | 71.079 | 1.533 | 2.153 | 55.844 |
| 0.18 | 33 (6.4) | 2.166 | 2.515 | 2.204 | 2.366 | 3.126 |
| 0.18 | 65 (12.7) | 2.146 | 3.284 | 2.244 | 2.361 | 4.316 |
| 0.18 | 129 (25.2) | 2.043 | 5.202 | 2.307 | 2.345 | 6.632 |
| 0.18 | 257 (50.2) | 1.787 | 3.194 | 2.302 | 2.318 | 12.414 |
| 0.18 | **513 (100.2)** | **1.459** | **0.324** | 2.123 | 2.266 | 26.155 |
| 0.18 | 1025 (200.2) | 1.470 | 71.217 | 1.503 | 2.153 | 55.844 |
| 0.2 | 33 (6.4) | 2.076 | 2.538 | 2.246 | 2.366 | 3.133 |
| 0.2 | 65 (12.7) | 2.060 | 3.308 | 2.263 | 2.361 | 4.414 |
| 0.2 | 129 (25.2) | 1.975 | 5.217 | 2.310 | 2.348 | 6.618 |
| 0.2 | 257 (50.2) | 1.755 | 3.261 | 2.290 | 2.318 | 12.448 |
| 0.2 | **513 (100.2)** | **1.489** | **0.378** | 2.089 | 2.266 | 23.892 |
| 0.2 | 1025 (200.2) | 1.499 | 71.217 | 1.457 | 2.153 | 55.844 |

**Supplementary Table S2: Continuous simulation benchmarking across window sizes and noise levels.** For each noise level, multiple candidate window sizes are evaluated, following an odd-length sequence of the form $2^n + 1$, ranging from 33 to 1025 samples, corresponding to window durations from 6.45 to 200.20 ms. The NeuroTD quality score is used to select the optimal window; the minimum quality score is highlighted in bold, and the corresponding window size and NeuroTD RMSE (alignment error, in ms) are also shown in bold. For each method (NeuroTD, DTW, CTW, and FFT), the minimum RMSE across window sizes within a given noise level is underlined, indicating oracle performance. For eight out of ten nonzero noise levels, the window selected by the NeuroTD quality score matches the RMSE-optimal window, and in the remaining cases yields closely comparable RMSE values, indicating that quality-based selection reliably identifies near-optimal windows without access to ground-truth delay information.

| Noise | Window (samples, ms) | Quality | NeuroTD | DTW | CTW | FFT |
|---|---|---|---|---|---|---|
| 0.000 | 33 (6.4) | 0.434 | 0.047 | 0.319 | 1.296 | 1.337 |
| 0.000 | 65 (12.7) | 0.384 | 0.043 | 0.184 | 1.298 | 1.339 |
| 0.000 | 129 (25.2) | 0.369 | 0.057 | 0.106 | 1.303 | 1.345 |
| 0.000 | **257 (50.2)** | **0.339** | **0.135** | 0.090 | 1.314 | 1.356 |
| 0.000 | 513 (100.2) | 0.385 | 0.273 | 0.202 | 1.332 | 1.375 |
| 0.000 | 1025 (200.2) | 0.602 | 0.524 | 0.541 | 1.332 | 1.386 |
| 0.02 | 33 (6.4) | 2.311 | 1.198 | 1.166 | 1.281 | 2.032 |
| 0.02 | 65 (12.7) | 2.032 | 1.528 | 1.151 | 1.284 | 3.047 |
| 0.02 | 129 (25.2) | 1.540 | 1.908 | 1.132 | 1.289 | 4.731 |
| 0.02 | 257 (50.2) | 0.993 | 1.131 | 0.882 | 1.300 | 6.451 |
| 0.02 | **513 (100.2)** | **0.692** | **0.293** | 0.694 | 1.318 | 11.609 |
| 0.02 | 1025 (200.2) | 0.749 | 0.535 | 0.774 | 1.317 | 25.895 |
| 0.04 | 33 (6.4) | 2.443 | 1.399 | 1.263 | 1.281 | 2.088 |
| 0.04 | 65 (12.7) | 2.236 | 1.864 | 1.287 | 1.283 | 3.333 |
| 0.04 | 129 (25.2) | 1.866 | 2.667 | 1.283 | 1.289 | 5.679 |
| 0.04 | 257 (50.2) | 1.339 | 1.996 | 1.579 | 1.300 | 9.265 |
| 0.04 | 513 (100.2) | 0.940 | 0.428 | 1.585 | 1.318 | 16.649 |
| 0.04 | **1025 (200.2)** | **0.935** | **0.561** | 1.291 | 1.316 | 39.138 |
| 0.06 | 33 (6.4) | 2.441 | 1.518 | 1.300 | 1.281 | 2.131 |
| 0.06 | 65 (12.7) | 2.291 | 1.994 | 1.362 | 1.284 | 3.481 |
| 0.06 | 129 (25.2) | 2.002 | 3.272 | 1.501 | 1.289 | 5.997 |
| 0.06 | 257 (50.2) | 1.553 | 2.843 | 1.809 | 1.300 | 10.724 |
| 0.06 | 513 (100.2) | 1.123 | 0.514 | 2.187 | 1.318 | 21.199 |
| 0.06 | **1025 (200.2)** | **1.094** | **0.577** | 1.792 | 1.316 | 42.783 |
| 0.08 | 33 (6.4) | 2.419 | 1.571 | 1.320 | 1.281 | 2.159 |
| 0.08 | 65 (12.7) | 2.302 | 2.009 | 1.402 | 1.284 | 3.553 |
| 0.08 | 129 (25.2) | 2.065 | 3.734 | 1.614 | 1.289 | 6.228 |
| 0.08 | 257 (50.2) | 1.679 | 3.470 | 2.079 | 1.300 | 11.210 |
| 0.08 | 513 (100.2) | 1.259 | 0.586 | 2.552 | 1.317 | 23.442 |
| 0.08 | **1025 (200.2)** | **1.223** | **0.584** | 2.325 | 1.316 | 45.597 |
| 0.1 | 33 (6.4) | 2.380 | 1.598 | 1.343 | 1.281 | 2.187 |
| 0.1 | 65 (12.7) | 2.288 | 2.064 | 1.452 | 1.283 | 3.608 |
| 0.1 | 129 (25.2) | 2.086 | 4.003 | 1.673 | 1.289 | 6.429 |
| 0.1 | 257 (50.2) | 1.760 | 4.385 | 2.297 | 1.300 | 11.709 |
| 0.1 | 513 (100.2) | 1.368 | 0.679 | 2.768 | 1.317 | 24.663 |
| 0.1 | **1025 (200.2)** | **1.328** | **0.611** | 2.363 | 1.316 | 47.332 |
| 0.12 | 33 (6.4) | 2.333 | 1.626 | 1.365 | 1.281 | 2.191 |
| 0.12 | 65 (12.7) | 2.258 | 2.105 | 1.470 | 1.283 | 3.666 |
| 0.12 | 129 (25.2) | 2.085 | 4.084 | 1.728 | 1.289 | 6.550 |
| 0.12 | 257 (50.2) | 1.802 | 4.748 | 2.348 | 1.300 | 12.047 |
| 0.12 | 513 (100.2) | 1.449 | 0.776 | 2.838 | 1.317 | 25.461 |
| 0.12 | **1025 (200.2)** | **1.409** | **0.639** | 2.343 | 1.316 | 48.935 |
| 0.14 | 33 (6.4) | 2.282 | 1.649 | 1.377 | 1.281 | 2.206 |
| 0.14 | 65 (12.7) | 2.216 | 2.144 | 1.490 | 1.283 | 3.727 |
| 0.14 | 129 (25.2) | 2.071 | 4.155 | 1.773 | 1.288 | 6.676 |
| 0.14 | 257 (50.2) | 1.834 | 5.243 | 2.296 | 1.300 | 12.320 |
| 0.14 | 513 (100.2) | 1.509 | 0.874 | 2.797 | 1.317 | 26.055 |
| 0.14 | **1025 (200.2)** | **1.472** | **0.654** | 2.499 | 1.315 | 50.076 |
| 0.16 | 33 (6.4) | 2.230 | 1.668 | 1.390 | 1.281 | 2.219 |
| 0.16 | 65 (12.7) | 2.172 | 2.173 | 1.503 | 1.283 | 3.755 |
| 0.16 | 129 (25.2) | 2.043 | 4.213 | 1.798 | 1.289 | 6.794 |
| 0.16 | 257 (50.2) | 1.838 | 5.538 | 2.296 | 1.300 | 12.438 |
| 0.16 | 513 (100.2) | 1.555 | 1.010 | 2.754 | 1.317 | 26.344 |
| 0.16 | **1025 (200.2)** | **1.519** | **0.656** | 2.319 | 1.315 | 50.899 |
| 0.18 | 33 (6.4) | 2.179 | 1.688 | 1.409 | 1.281 | 2.236 |
| 0.18 | 65 (12.7) | 2.127 | 2.200 | 1.518 | 1.284 | 3.776 |
| 0.18 | 129 (25.2) | 2.011 | 4.187 | 1.807 | 1.289 | 6.867 |
| 0.18 | 257 (50.2) | 1.833 | 5.825 | 2.352 | 1.300 | 12.601 |
| 0.18 | 513 (100.2) | 1.585 | 1.135 | 2.802 | 1.317 | 26.763 |
| 0.18 | **1025 (200.2)** | **1.554** | **0.660** | 2.540 | 1.316 | 51.876 |
| 0.2 | 33 (6.4) | 2.128 | 1.705 | 1.426 | 1.280 | 2.243 |
| 0.2 | 65 (12.7) | 2.082 | 2.232 | 1.535 | 1.284 | 3.809 |
| 0.2 | 129 (25.2) | 1.975 | 4.201 | 1.839 | 1.289 | 6.955 |
| 0.2 | 257 (50.2) | 1.822 | 6.084 | 2.375 | 1.300 | 12.705 |
| 0.2 | 513 (100.2) | 1.607 | 1.237 | 2.695 | 1.317 | 27.016 |
| 0.2 | **1025 (200.2)** | **1.581** | **0.645** | 2.364 | 1.316 | 52.295 |

### Supplementary Figures

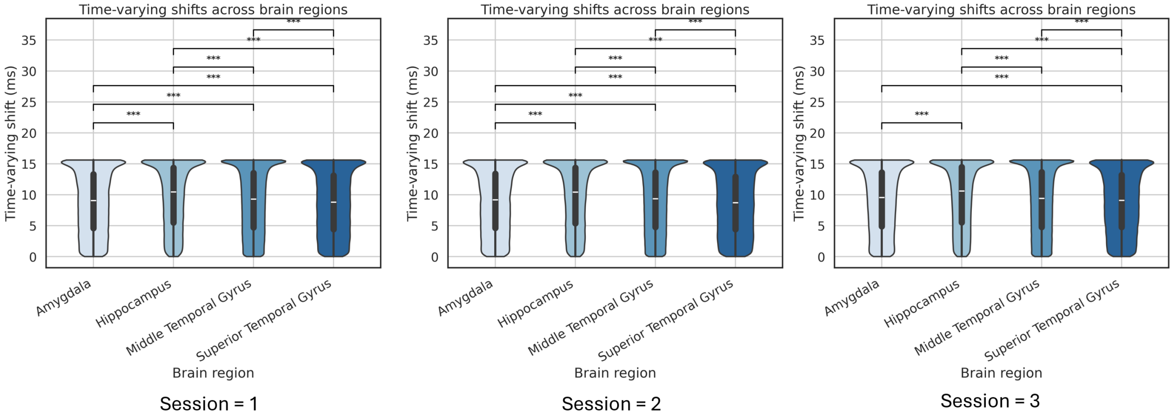


**Supplementary Figure S1: Consistency of time-varying neural delays across experimental sessions.** Violin plots show the distribution of time-varying delays stratified by anatomical brain region for each session of a human subject, analyzed independently. Across all sessions, the hippocampus consistently shows the highest median delay, while the superior temporal gyrus shows the lowest. The overall ranking and statistical differences among regions are robust to session-to-session variability, supporting the findings in the main figure 3c that these delay patterns are reproducible across experimental repeats.

### Supplementary Data

**Supplementary Data S1:** Top 200 genes ranked by the absolute value of their correlation with ISTD scores. The dataset is provided as a `.csv` file containing gene names and corresponding signed correlation coefficients, and was used for downstream enrichment analysis.

The file is available as `Supplementary_Data_S1_genes_top200.csv`.
